# Fluorescent protein-based reporters reveal stress response of intracellular *Salmonella enterica* on single cell level

**DOI:** 10.1101/2020.07.20.213272

**Authors:** Marc Schulte, Katharina Olschewski, Michael Hensel

## Abstract

Intracellular bacteria such as *Salmonella enterica* are confronted with a broad array of defense mechanisms of their mammalian host cells. The ability to sense host cell-imposed damages, and to mount efficient stress responses are crucial for survival and proliferation of intracellular pathogens. The various combinations of host defense mechanisms acting on intracellular bacteria and their individual response also explain the occurrence of distinct subpopulations of intracellular *S. enterica* such as dormant or persisting, slowly or rapidly replicating cells. Here we describe a set of fluorescence protein (FP)-based reporter strains that were used to monitor the expression of cytoplasmic or periplasmic stress response systems on a single cell level. This is mediated by a fast maturing FP as reporter for induction of stress response genes. We evaluated slower maturing FPs for a second function, i.e. the analyses of the status of intracellular proliferation of pathogens. The combination of two FPs allows, on a single cell level, the interrogation of stress response and intracellular proliferation. Application of these reporters to *S. enterica* allowed us to detect and quantify distinct intracellular subpopulations with different levels of stress response and proliferation.

**Importance:** Sensing of, and responding to host-mediated damages are important defensive virulence traits of bacterial pathogens. Intracellular pathogens such as *Salmonella enterica* are exposed to various types of antimicrobial host cell defenses that impose, among other, periplasmic and cytosolic stresses. Intracellular *S. enterica* form distinct subpopulations that differ in proliferation rate, metabolic activity and persister formation. Here we deploy fluorescence protein-based reporter strains to monitor, on a single cell level, the response of intracellular *S. enterica* to periplasmic or cytoplasmic stress. A second fluorescent protein reports the biosynthetic capacity of individual intracellular *S. enterica*. The dual fluorescence reporters can be deployed to characterize by flow cytometry phenotypically diverse subpopulations and stress responses in intracellular bacteria.

## Introduction

The ability to sense environmental cues and to respond to potentially detrimental factors is of vital importance for bacterial cells. In addition to various common stress factors, pathogenic and facultative intracellular bacteria such as *Salmonella enterica* serovar Typhimurium (STM) face various defense mechanisms of the host that normally result in clearance of bacteria that breach borders within a host. The proper response to environmental or host-imposed damages thus is an important part of the defensive virulence factors. As attack of host cells against intracellular pathogens is multifactorial and acting on various bacterial structures, stress responses have to perform efficiently in order to mount protection and repair that is coordinated in space and time. Gram-negative bacteria possess an array of stress response systems (SRS) that can sense harmful conditions and damages to the cell envelope, or to periplasmic or cytosolic components. Periplasmic stress often results in the accumulation of misfolded proteins that cannot be properly assembled into polymeric surface structures such as fimbriae, or inserted into the outer membrane. A common response to the accumulation of misfolded or otherwise damaged proteins in the periplasm is activation of proteolytic degradation (reviewed in 1). Here, periplasmic protease HtrA, a.k.a. DegP has a key role as multifunctional protein quality control factor. HtrA functions both as proteolytic enzyme and chaperone with broad substrate specificity, but only targeting misfolded and/or mislocalized proteins. Such proteins are recognized by the specific PDZ domain 1 found in proteins important for signal transduction and proteolytic activity. HtrA has a unique molecular architecture, i.e. a serine protease domain and two PDZ domains per monomer. Their activity is regulated by the conserved mechanism by oligomerization. To support refolding of denatured proteins, peptidyl-prolyl isomerases such as FkpA are present in the periplasm. Their main mode of action is the *cis-trans* isomerization of peptidyl-prolyl bonds. It is a crucial step in protein folding because both *cis* and *trans* peptide bonds are found in prolines of native proteins. However, the isomeric state of prolines in each protein is characteristic and important for proper protein function. FkpA belongs to the FKBP family because of its binding to FK506. In addition, FkpA acts as chaperone with activity independent of its peptidyl-prolyl isomerase activity (2). Expression of factors responding to periplasmic stress is coordinately regulated by DegS and the sigma factor E pathway, the two-component system BaeRS, and two-component system CpxAR with its periplasmic accessory component CpxP (reviewed in 3).

Another form of damage is the loss of turgor resulting from hyperosmotic environments, or damage of the cytoplasmic membrane barrier. As one form of counteraction, the periplasmic trehalase TreA is expressed that catalyzes hydrolysis of osmo-protectant trehalose into two glucose molecules (reviewed in 4).

The cytosolic stress response is required, among others, for repair of oxidized cysteine residues in cytosolic proteins that is achieved by functionally partially redundant thioredoxins (Trx) and glutaredoxins (Grx) (reviewed in 5, 6). Trx reduces a disulfide bond in an oxidized substrate protein. The reaction starts with a nucleophilic attack on the disulfide bond carried out by the amino-terminal cysteine of the specific CXXC motif of Trx. A mixed-disulfide intermediate complex is formed subsequently that undergoes further reduction by another nucleophilic attack carried out by the carboxy-terminal cysteine of the specific CXXC motif. This results in release of a reduced and ‘repaired’ substrate protein. Oxidized Trx is further reduced by Trx reductase TrxR, which is reduced by NADPH + H^+^. The mode of action of Grx is comparable to Trx. However, after the release of the reduced substrate protein and the oxidized Grx, the oxidized Grx is further reduced by two glutathione (GSH) molecules. Oxidized glutathiones are then reduced to GSH by a glutathione reductase using NADPH + H^+^ (5).

MsrA and BisC are required to repair oxidized methionine residues. MsrA acts in concert with MsrB to repair reactive oxygen species (ROS)-induced oxidative damages. The catalytic mechanism operates by a so-called cysteine-based three-step mechanism. (i) A complex of MsrA or MsrB with methionine sulfoxide is formed. Afterwards, the catalytic cysteine attacks the sulfoxide by a nucleophilic attack. A sulfenic acid intermediate is formed and the ‘repaired’ and reduced substrate protein is released. (ii) The recycling cysteine of MsrA/B attacks the sulfenic acid intermediate which creates an intramolecular disulfide bond. (iii) Trx is used to reduce the intramolecular disulfide bond. The mode of action of BisC is comparable to MsrA and MsrA, however, whereas MsrAB is able to reduce both free Met-O and Met-O in proteins, BisC is only able to reduce free Met-O (7–9).

Next to the defensive mechanisms required to survive within the host, STM has evolved various virulence mechanisms to actively manipulate the host. The intracellular lifestyle of STM within the *Salmonella*-containing vacuole (SCV) in host cells is critically dependent on the function of the type III secretion system (T3SS) encoded by *Salmonella* pathogenicity island 2 (SPI2) (10). Effector proteins translocated by the SPI2-T3SS remodel the host cell endosomal system, resulting in the formation of an extensive network of interconnected membrane vesicles termed *Salmonella*-induced filaments (SIF). This action of the SPI2-T3SS has been linked to the nutrition of intracellular STM, the avoidance of host cell defense mechanism, and reduction of exposure to stress factors (11–13).

Although STM is equipped with this range of defensive and offensive virulence factors and able to deploy these, the individual fate of intracellular STM is surprisingly heterogenous (14). While a subset of STM cells initiate rapid intracellular proliferation, another subset remains non-replicating or dormant, yet another subset is killed by antimicrobial defenses of the host cell. If the divergent intracellular fate of STM is a consequence of the ability to respond to stress factors remains to be determined.

Analyses of mutant strains deficient in SRS have been performed, but did not reveal the role of stress response on single cell level. We aimed to generate a reporter system for single cell analyses of response of STM to stress within host cells, and as well allows quantification of bacterial proliferation. Here we introduce a dual fluorescence reporter system that enables, for single intracellular STM cells, analysis of activation of specific stress responses and intracellular proliferation.

## Results

### *Design of fluorescent protein-based reporters for stress response of intracellular* Salmonella enterica *on single cell level*

We aimed to analyze, on the level of single bacterial cells, the sensing and response of stress by intracellular STM. For this, the basic design of dual color fluorescence reporters was applied (15) as shown in **Fig. 1**. For single cell analyses, the reporter should allow bacterial detection by flow cytometry (FC), as well as by epi-fluorescence microscopy. After selection of the fluorescent protein (FP) as described below, the plasmid vectors consisted of an expression cassette for DsRed version DsRed T3_S4T (16) under control of a synthetic constitutive promoter P_EM7_, and sfGFP under control of the regulated promoter. DsRed expression was homogeneous among individual cells of STM under culture conditions and allowed detection of 97.0–99.6 % of the all bacteria.

**Fig. 1.**
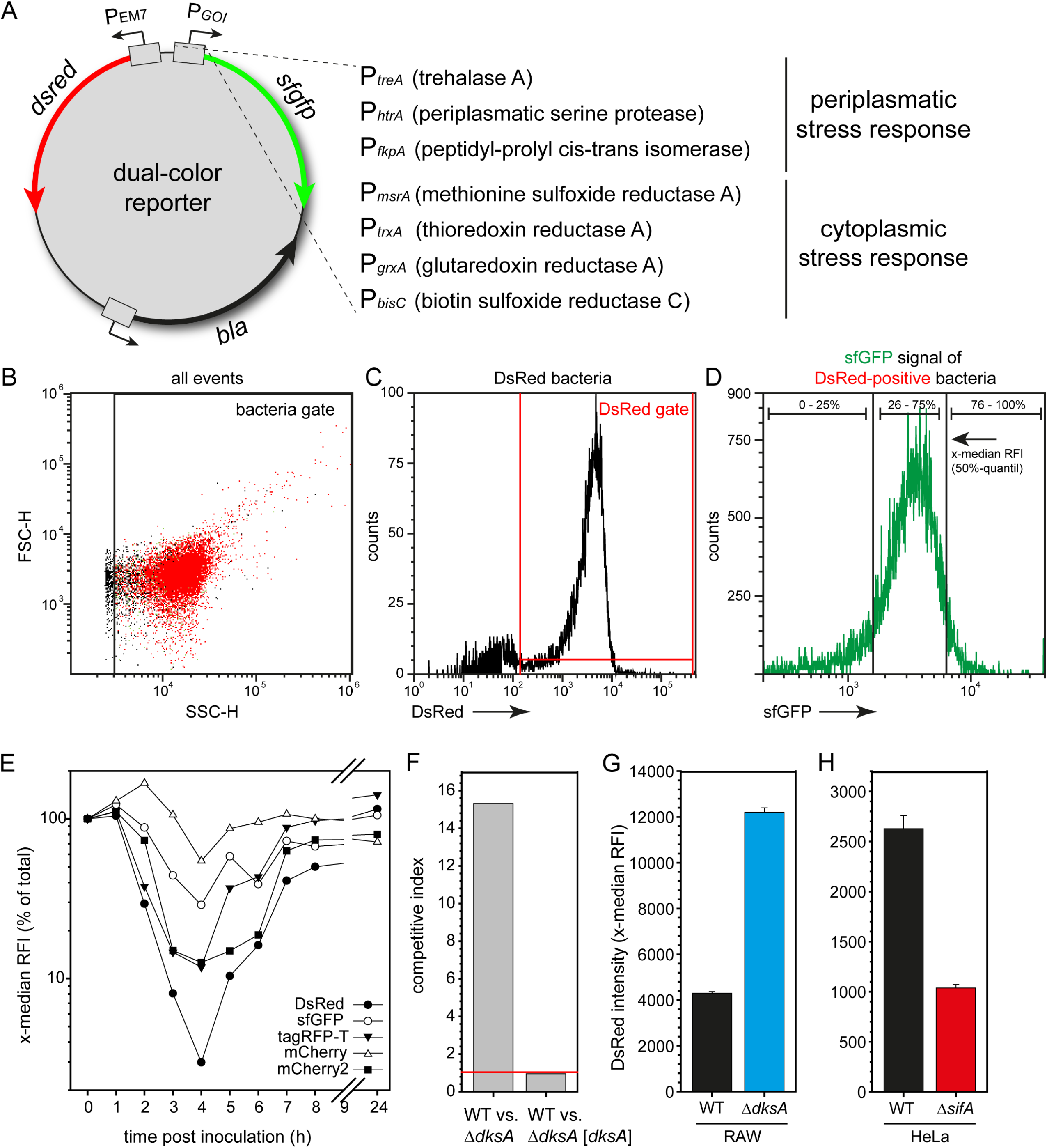
Design of dual color stress reporters. A) Dual color reporter plasmids encode DsRed under control of the constitutive promoter P_EM7_. sfGFP is encoded under control of a stress-induced promoter of the gene of interest (GOI), indicated as P_GOI_. In this study, three promoters of genes involved in response to periplasmic stress (PSR) were analyzed i.e. *treA*, *htrA*, and *fkpA*. The promoters of *msrA*, *trxA*, *grxA*, and *bisC* were used to monitor cytoplasmic stress response (CSR). The constitutive DsRed expression allowed identification of bacteria within host cells, or after release from host cells. The levels of sfGFP fluorescence are anticipated to correlate to levels of specific stress sensed by STM. B, C, D) Gating approach for flow cytometry (FC) analyses of STM harboring dual color reporter. The FSC/SSC was set of identify bacteria-sized particles in the bacteria gate (B), the DsRed fluorescent events in the bacteria gate were gated (C), and the sfGFP intensity of DsRed-positive bacteria was quantified (D). E) STM WT harboring plasmids with constitutive expression of either *dsred*, *sfgfp*, tag-*rfpT*, *mCherry* or *mCherry2* was grown o/n in LB medium, diluted to an OD_600_ of 0.05 in fresh LB and incubated at 37°C. In intervals of 1 h and after culturing for 24 h, samples were taken and subjected to FC to measure the constitutive fluorescence intensity. The x-median relative fluorescence intensity (RFI) always represents the constitutive fluorescence signal of the entire bacterial population. F) RAW264.7 macrophages were coinfected with STM WT and Δ*dksA* harboring a resistance marker or a complementation plasmid. X-fold replication after 16 h p.i. was determined and the competitive index was calculated. Competitive index shows high replication defect of Δ*dksA* compared to STM WT. Deletion phenotype is able to be recovered by introduction of a complementation plasmid into Δ*dksA*. G, H) STM WT, Δ*dksA,* or Δ*sifA* strains, each harboring dual color fluorescence reporter for *msrA* were used to infect RAW264.7 macrophages (G) or HeLa cells (H). Infected cells were lysed 24 h (G) or 8 h p.i. (H). Liberated STM were recovered, fixed and subjected to FC to quantify the DsRed intensity per STM cells. Means and standard deviations of triplicate samples are shown, and the experiments shown is representative for three replicates

Because various FP have divergent maturation times, we analyzed the correlation of growth and fluorescence dilution for a set of commonly used FP. For this, STM WT harbored plasmids for constitutive expression of DsRed, tagRFP-T, mCherry, mCherry2, or sfGFP under control of P_EM7_. The constitutive expression of the various FP did not affect growth yield or growth rate in LB (**Fig. S 1**A). Despite the reported differences in toxicity of some FP, all FP were sufficiently well tolerated by STM at the expression levels used for the dual color reporters.

We analyzed the effect of the bacterial growth phase on dilution of DsRed (**Fig. S 1**B). STM WT harboring reporter [P_EM7_::*dsred* P_*cypD*_::*sfgfp frameshift*] (red control) was grown in LB medium and samples were collected in hourly intervals for quantification of DsRed fluorescence levels by FC. In STM entering the log phase, a steep drop of DsRed fluorescence was observed. The x-median RFI was 27.7-fold lower in cells at 4 h of subculture, i.e. transition from late log phase to early stationary phase. After entering stationary phase and reaching a constant cell density, a continuous increase of DsRed fluorescence per cell was determined.

Analyses of relative fluorescence intensities (RFI) of various FP during time of culture in LB (**Fig. 1**E) demonstrated that mCherry and sfGFP showed rather constant FP intensities, with decrease to 54.5 % and 28.9 % of initial x-median RFI, respectively. The red FP tagRFP-T and mCherry2 revealed more pronounced decrease to 11.8 % and 12.9 % of initial intensity at 4 h of subculture. While the decrease of x-median RFI in log phase culture was comparable for both FP, tagRFP-T showed more rapid increase in RFI than mCherry2 in STM entering the stationary phase. Of the FP analyzed, DsRed showed the strongest growth phase-dependent decrease of x-median RFI to a minimum of 3.0 % of initial fluorescence (**Fig. 1**E).

For further correlation of growth kinetics to DsRed dilution, we cultured STM in media that lead to different growth rates (**Fig. S 1**CD). Growth in synthetic PCN (Phosphate, Carbon, Nitrogen) medium resulted in culture yield comparable to LB medium, but altered growth kinetics. Lowest growth rate was observed in PCN with maltose as C-source (0.66), moderate growth rate in PCN with glucose (0.98), and highest growth rate in LB (1.33) (**Fig. S 1**C). Quantification of DsRed fluorescence again showed a sharp decrease in RFI during culture in LB with a minimum at 4 h of subculture 3.6 % of the initial DsRed intensity of the inoculum (**Fig. S 1**D), while the decrease was less pronounced for PCN glucose cultures, with a delayed minimum at 6 h of subculture at transition to stat. phase (26.6 % of inoculum RFI). The slow growth in PCN maltose correlated with rather constant DsRed fluorescence levels during subculture. Here, minimal fluorescence was detected at 7 h of subculture at transition to stat. phase with 74.2 % of inoculum RFI. These data confirm that FP dilution correlates with STM growth rate and may serve as proxy for the level of intracellular proliferation and growth state of STM.

We conclude that the long maturation time of DsRed, and the dilution by growing cultures renders this FP as suitable reporter for the evaluation of growth kinetics of STM. Beside the primary use for detection of intracellular STM, constitutive DsRed expression may serve as an additional reporter for the growth state of STM. Despite the large range of DsRed fluorescence intensities in the various phases of culture, the DsRed fluorescence levels were sufficiently high to detect STM in FC by virtue of red fluorescence and allowed discrimination from bacteria-sized debris in crude lysates of infected host cells.

The second gene for a FP, here the bright and fast maturing sfGFP, was under control of a stress-induced promoter. We selected a set of promoters of genes known to be involved in periplasmic stress response (PSR), i.e. *treA*, *htrA*, and *fkpA*, and in cytosolic stress response (CSR), i.e. *msrA*, *trxA*, *grxA*, and *bisC*. The modular design of the reporter plasmids facilitates exchange of promoter elements and genes for FP to meet specific experimental demands.

The dual color reporter P_EM7_::*dsred* P_*msrA*_::*sfgfp* was analyzed in STM WT and Δ*dksA* background (**Fig. 1**FG). Intracellular proliferation of the Δ*dksA* strain was highly attenuated in RAW264.7 macrophages, and a competitive index of 15.3 was determined for intracellular proliferation of STM WT vs. Δ*dksA*, complementation of Δ*dksA* restored the competitive index to 0.95-fold (**Fig. 1**F). Analyses of DsRed fluorescence in STM recovered from RAW264.7 cells 8 (**Fig. 1**G) revealed highly increased red fluorescence per bacterial cell for STM Δ*dksA* (12,297 x-median RFI) compared to STM WT (4,305 x-median RFI). STM deficient in *sifA* frequently lose SCV integrity, are released into the host cell cytosol and show rapid proliferation (hyper-replication) in epithelial cells (17, 18). Analyses of STM after infection of HeLa cells indicated a strong decrease of DsRed fluorescence for the Δ*sifA* strain compared to STM WT (**Fig. 1**H). This decrease was also obvious in time lapse microscopy of infected HeLa with hyper-replicating STM (data not shown) and is in line with the dilution of DsRed in rapidly replicating STM. These results indicate that DsRed fluorescence levels report growth levels of intracellular STM.

### Dual color reporters faithfully report specific stress exposure

As a functional control for the performance of the stress reporter plasmids, we used *in vitro* cultures of STM with exposure to defined stressors (**Fig. 2 Error! Reference source not found.**, **Fig. S 2**). We determined the response of the reporter P_*msrA*_::*sfgfp* to various concentrations of H_2_O_2_ or methyl viologen (MV) (**Fig. S 2**A), and reporter P_*htrA*_::*sfgfp* to various concentrations of DTT or polymyxin B (**Fig. S 2**B). Furthermore, the growth phase-dependent expression of the various stress reporters was analyzed (**Fig. S 2**C). The analyses revealed that P_*treA*_ was highly induced in stationary cultures (5.5-fold), while P_*htrA*_ and P_*msrA*_ showed about 2.4-fold increased induction in stationary phase cells. The other reporters showed no increased induction in stationary phase (**Fig. S 2**D).

**Fig. 2.**
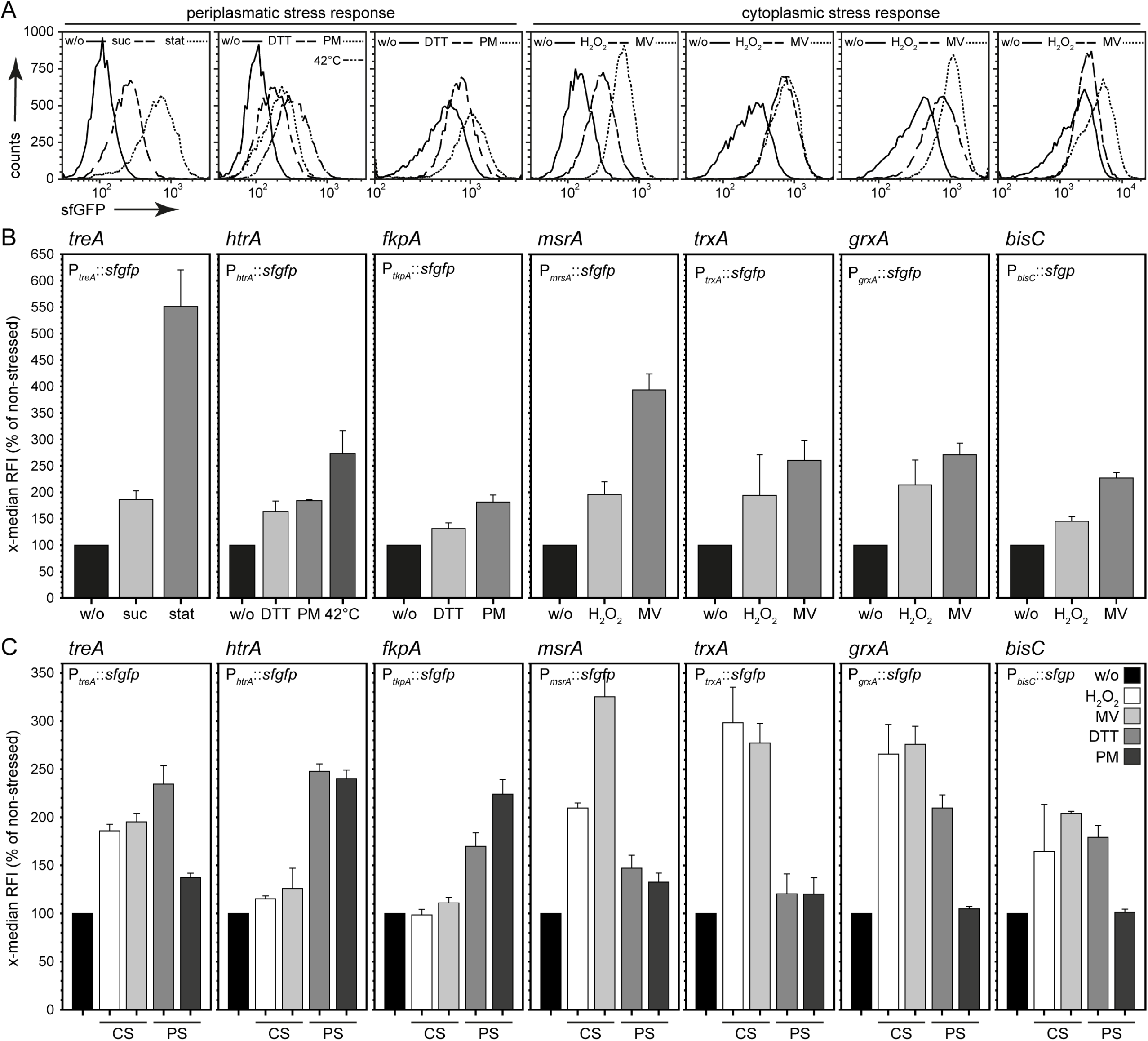
*In vitro* induction of stress reporter using various stressors. STM WT harboring dual color fluorescence reporter for *treA*, *htrA* or *fkpA* representing the PSR, and *msrA*, *trxA*, *grxA* or *bisC* representing the CSR was grown in PCN minimal medium o/n and diluted in fresh PCN medium for further subculture. After 3 h of subculture bacteria were stressed using 1 mM H_2_O_2_, 0.3 mM methyl viologen (MV), 3 mM DTT, 10 μg × ml^−1^ polymyxin B (PM), 1 M sucrose (suc), or 42°C heat shock for 3 h. For additional induction of P_*treA*_, a stationary culture was used (stat). As control, no stressor was added (w/o). Subsequently, samples were taken, fixed and subjected to FC. The x-median was normalized to non-stressed STM and represents the induced sfGFP signal of the entire DsRed-positive bacterial population (B), and the respective histograms are shown in A). C) Cross-induction of CS reporter (*msrA*, *trxA*, *grxA* or *bisC*) responding to PS (imposed by DTT or PM), and PS reporter (*treA*, *htrA* or *fkpA*) responding to CS (imposed by H_2_O_2_ or MV) were analyzed. Means and standard deviations of three biological replicates are shown.

Based on these data, STM was exposed to 1 mM H_2_O_2_ or 0.3 mM methyl viologen in the late log phase of subculture in chemically defined media to induce cytosolic stress. For periplasmic stress induction, STM was exposed to 3 mM DTT, or 10 μg × ml^−1^ polymyxin B (PM). Furthermore, osmotic stress was imposed by 1 M sucrose (suc), or heat shock by shift of cultures to 42 °C for 3 h. For additional induction of *treA*, culture growth to stationary phase was used (stat). The levels of sfGFP expression in non-stressed and stress-exposed STM were determined by FC. As indicated in **Fig. 2**A and B, each of the stress reporters showed a specific induction by stressor addition, or stress exposure. However, the levels of induction varied between the reporters. For CSR, we found that the P_*msrA*_ fusion resulted in the highest induction (3.9-fold) of sfGFP fluorescence, while 5.5-fold induction was observed for PSR sensor P_*treA*_ in stationary cultures. In addition, stress reporters were checked for cross-induction to other stressors than what they should be reacting to (**Fig. 2**C). For that, PSR reporters were tested for response to CS, and *vice versa*. Concentrations and application of stressors was performed as described above and the levels of sfGFP expression were determined by FC. Reporters for PSR (P_*htrA*_ and P_*fkpA*_) only showed induction upon exposure to periplasmic stressors DTT and PM, whereas no induction could be observed when H_2_O_2_ or MV was added, indicating specificity of the reporters. The P_*treA*_::*sfgfp* fusion was induced when H_2_O_2_, MV or DTT were added, but only low induction was observed after PM treatment. The reporters for the CSR (P_*msrA*_, P_*trxA*_, P_*grxA*_, and P_*bisC*_) only responded to CS triggered by H_2_O_2_ or MV. Increased induction after addition of the PS inductor PM was not observed. However, after stressing with DTT, slight induction of P_*msrA*_ and P_*trxA*_, and higher induction of P_*grxA*_ and P_*bisC*_ was detected. Thus, also the reporters P_*msrA*_ and P_*trxA*_ for CRS showed specificity to CS. A low degree of cross-induction was observed for CSR reporters P_*grxA*_ and P_*bisC*_ after addition of the periplasmic stressor PM.

### *Response of reporters to stress exposure in intracellular* Salmonella

We next analyzed the response of stress reporters in intracellular STM. STM WT harboring the various reporters for PSR or CSR were used to infect the human epithelial cell line HeLa, or the murine macrophages-like cell line RAW264.7 (**Fig. 3**). The intracellular bacteria were released from host cell 8 h p.i. and subjected to FC. For comparison, the bacterial inoculum was analyzed using the same settings for FC. STM WT harboring a plasmid for constitutive expression of DsRed, but lacking expression of sfGFP served as negative control for adjustment of FC gating. For all stress reporters investigated, we observed increased induction of sfGFP levels in intracellular STM (P_*fkpA*_::*sfgfp* and P_*mrsA*_::*sfgfp* were induced 7.32-fold and 7.34-fold, respectively, in HeLa cells, and 6.21-fold and 7.02-fold for P_*trxA*_::*sfgfp* and P_*grxA*_::*sfgfp*, respectively, in RAW264.7 cells. For most sensors, the overall sfGFP expression in intracellular bacteria was higher in RAW264.7 cell than in HeLa. For P_*bisC*_ and P_*fkpA*_, induction was comparable for intracellular bacteria in both cell lines. Since STM grown to stationary phase were used for RAW264.7 infection, the expression levels in the inoculum were higher than in late log phase cultures used for invasion of HeLa cells.

**Fig. 3.**
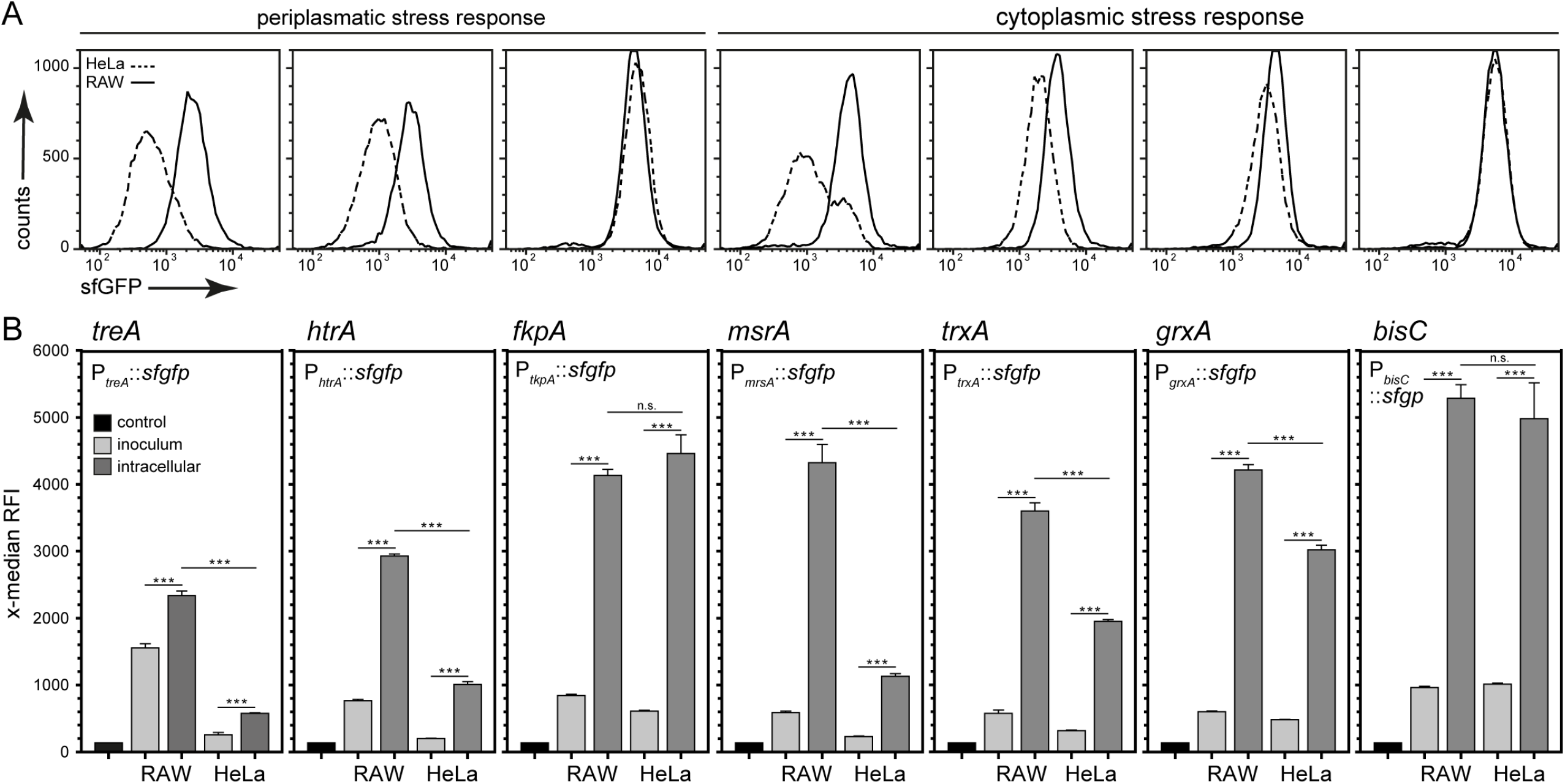
Intracellular periplasmic and cytoplasmic stress proteins are induced within murine macrophages and HeLa cells. STM WT harboring dual color fluorescence reporters for *treA*, *htrA* or *fkpA* representing the PSR, and *msrA*, *trxA*, *grxA,* or *bisC* representing the CSR were grown o/n in LB medium. RAW264.7 macrophages or HeLa cells were infected using reporter strains and lysed 8 h p.i. Liberated STM were recovered, fixed and subjected to FC. In addition, the inoculum for infection was fixed and subjected to FC. As non-induced negative control, the same dual color reporter backbone was used with a frame shift mutation in *sfgfp* (described before in (13). The x-median represents the induced sfGFP signal of the entire DsRed-positive intracellular bacterial population (B) and the respective histograms are shown in A). Means and standard deviations of one representative experiment are shown. Statistical analyses were accomplished by One Way ANOVA using SigmaPlot, and significance levels are indicated as follows: *, p < 0.05; **, p < 0.01; ***, p < 0.001; n.s., not significant.

We conclude that the various promoter fusions are suitable to report the stress exposure of intracellular STM.

### Role of detoxification and repair mechanisms

As pathogen adapted to life within host cells, STM efficiently deploys stress responses, detoxification and repair mechanisms to thrive in the vacuolar environment of the SCV. Therefore, we investigated the stress response induction of STM WT within γ-Interferon (IFNγ)-induced RAW264.7 macrophages at early time points post infection used P_*msrA*_ as representative for SR induction (**Fig. 4**A). We observed higher induction of P_*msrA*_::*sfgfp* reporter when analyzed in IFNγ-induced RAW264.7 macrophages. At 2, 4, 6, and 8 h p.i., the x-median RFI of sfGFP was significantly higher for STM in activated compared to resting RAW264.7 macrophages. Because the difference for P_*msrA*_::*sfgfp* induction was most pronounced at 2 h p.i. we next analyzed the induction of all stress reporter fusions at 2 h p.i. in resting and activated macrophages (**Fig. 4**B). At this time point, all reporter fusions apart from P_*treA*_::*sfgfp* showed an increased x-median RFI of sfGFP in STM in IFNγ-induced RAW264.7 compared to resting macrophages.

**Fig. 4.**
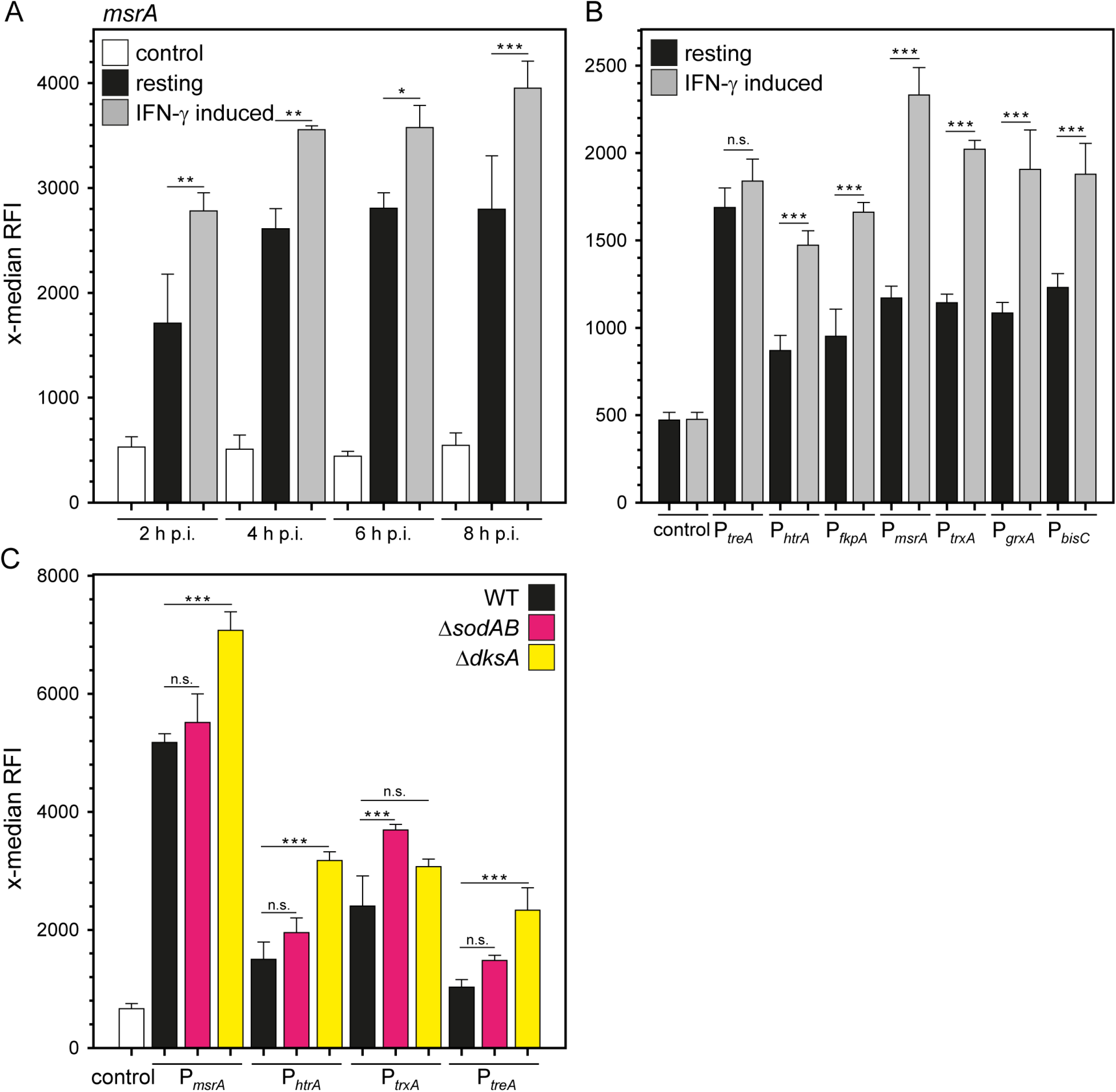
STM WT within interferon-γ-induced murine macrophages and STM mutant strains deficient in stress response show increased induction of stress reporters. A) STM WT harboring dual color reporter for *msrA* was grown o/n in LB medium. RAW264.7 macrophages were infected using reporter strains and lysed after various time points post infection. STM released from host cells were recovered, fixed and subjected to FC. As a non-induced negative control, the same dual color reporter backbone was used with a frame shift mutation in sfGFP. B) STM WT harboring dual color fluorescence reporters for *treA, htrA, fkpA, msrA, trxA, grxA* or *bisC* are shown. Infection and FC sample preparation was performed as described above. Host cells were lysed 2 h p.i. C) STM WT, Δ*sodAB* and Δ*dksA* harboring dual color fluorescence reporters for *msrA*, *htrA*, *trxA,* or *treA* are shown. Infection and FC sample preparation was performed as described above. Host cells were lysed 24 h p.i. The x-median represents the induced sfGFP signal of the entire DsRed-positive intracellular bacterial population. Means and standard deviations of one representative experiment are shown. Statistical analyses were performed as for **Fig. 3**.

In addition, we anticipated that in absence of functional detoxification and/or repair mechanisms, intracellular STM will be exposed to higher levels of stress, and in turn show increased induction of stress reporters. To test this model, we compared the expression of various stress reporters in STM WT to STM Δs*odAB* and STM Δ*dksA* strains. SodA and SodB are the cytosolic superoxide dismutases and essential for the detoxification of ROS (19). DksA is a transcriptional regulator and important in repair mechanisms for reactive nitrogen species (RNS)-mediated damages. Prior work demonstrated that Δs*odAB* and Δ*dksA* strains are hypersusceptible to ROS and RNS exposure, respectively, and attenuated in virulence (20).

For the stress reporters P_*msrA*_, P_*htrA*_, P_*trxA*_, and P_*treA*_, we observed higher induction in Δs*odAB* and Δ*dksA* strains (**Fig. 4**C). Strongest effects were determined in Δ*dksA* background with 2.12- and 2.27-fold increased x-median RFI for P_*htrA*_::*sfgfp* and P_*treA*_::*sfgfp* reporters, respectively. These data confirm that stress sensors report stress exposure of intracellular STM, and that functionality of detoxification and repair mechanisms affects expression levels.

### *Role of the SPI2-T3SS in ablation of stress in intracellular* Salmonella

In addition to SRS that are conserved in many bacteria, specific virulence factors also contribute to avoid exposure of intracellular bacteria to host defense mechanisms, and by this exposure to stress factors such as antimicrobial peptides, acidic pH, ROS and RNS, and nutritional deprivation. A role of the SPI2-T3SS and its effector proteins in converting the SCV into an environment permissive for survival and proliferation been proposed (11).

We used P_*htrA*_ and P_*msrA*_ as reporters representative for PSR and CSR, respectively. The reporters were introduced in STM WT, a *ssaV*-deficient strain unable to translocated SPI2-T3SS effector proteins, the *sifA*-deficient strain unable to induce endosomal remodeling, and the *sseF*-deficient strain that exhibit a reduced capacity in endosomal remodeling. While *ssaV* and *sifA* mutant strains are highly attenuated in virulence in murine models of systemic salmonellosis, the *sseF* mutant strain shows moderate attenuation (18, 21, 22).

Analyses of stress reporter induction for intracellular STM indicated strongest induction in the Δ*ssaV* background in RAW264.7 cells (**Fig. 5**, **Fig. S 3**, 1.6-fold and 1.4-fold for P_*msrA*_ and P_*htrA*_, respectively). These reporters showed also slightly higher induction in Δ*sifA* (only P_*msrA,*_, 1.22-fold) and Δ*sseF* background compared to STM WT (1.22-fold and 1.1-fold for P_*msrA*_ and P_*htrA*_, respectively). As shown in **Fig. 3**, induction of the reporters in HeLa cells was less pronounced. Comparison of induction in STM WT, Δ*ssaV*, Δ*sifA* and Δ*sseF* strains revealed only minute strain-specific differences. Expression of the P_*htrA*_ fusion in the Δ*sifA* background was significantly lower 0.75-fold) than for Δ*sseF* and Δ*ssaV* strains in RAW264.7 cells, and significantly lower compared to all other strains in HeLa cells. This reflects the release from the SCV for a part of the intracellular population of STM Δ*sifA* and relief from stress imposed within the SCV.

**Fig. 5.**
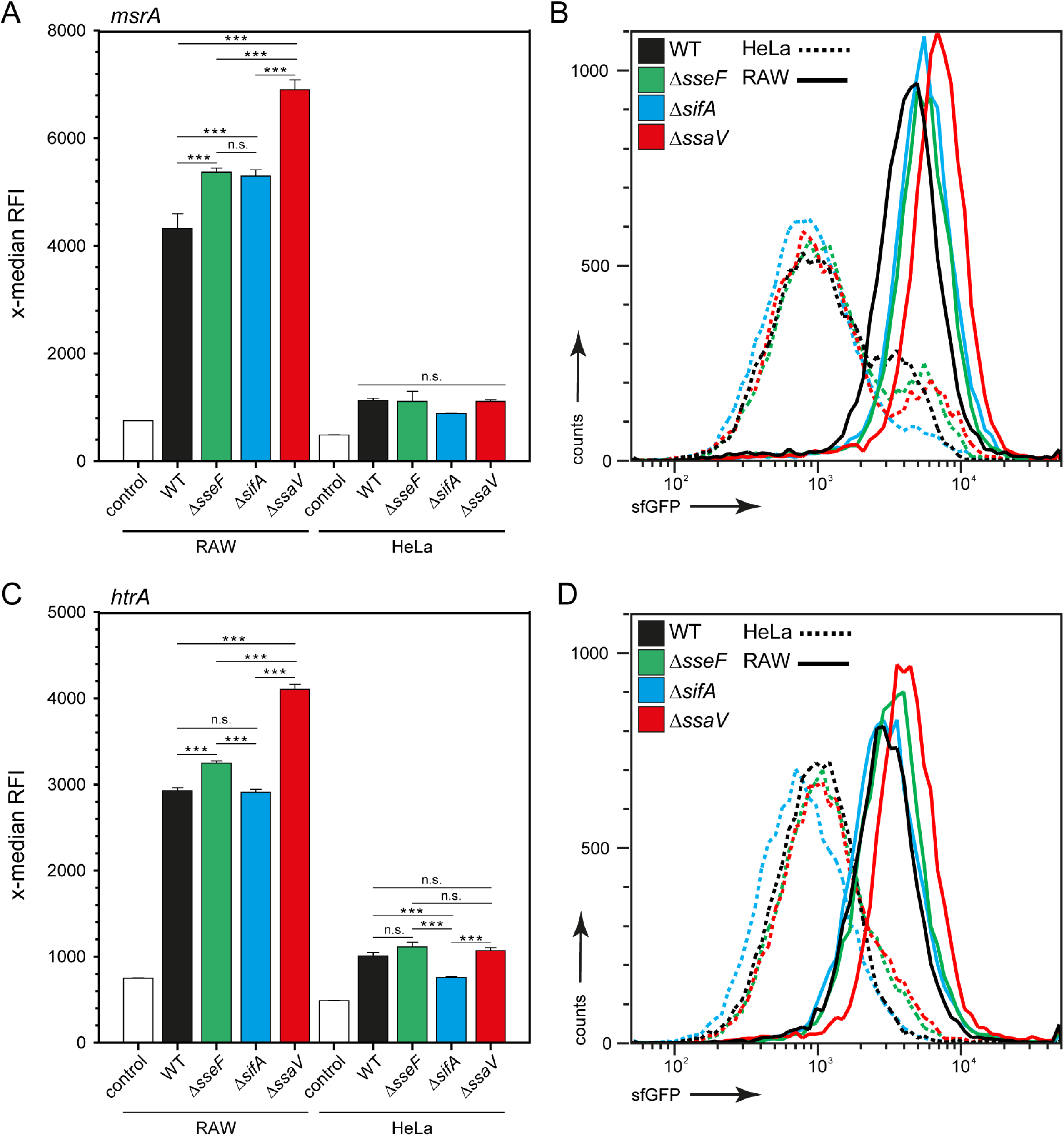
Higher stress induction of SPI2 mutant strains in macrophages, but not in HeLa cells. STM WT, Δ*sseF*, Δ*sifA* and Δ*ssaV* strains harboring dual color fluorescence reporter for *msrA* (A, B) and *htrA* (C, D) were grown o/n in LB medium. RAW264.7 macrophages or HeLa cells were infected using reporter strains and lysed 8 h p.i. Liberated STM were recovered, fixed and analyzed by FC. As non-induced negative control, the same dual color reporter backbone was used with a frame shift mutation in *sfgfp*. The x-median represents the induced sfGFP signal of the entire DsRed-positive intracellular bacterial population (A, C), and representative histograms are shown (B, D). Means and standard deviations of one representative experiment are shown. Statistical analyses were performed as for **Fig. 3**.

To test the coordination of stress response by intracellular STM, we analyzed the P_*msrA*_::*sfgfp* reporter in the background of a Δ*phoPQ* strain (**Fig. S 4**). This strain is highly attenuated due to the defect in a two-component system with global control of expression of stress response functions and virulence genes (reviewed in 23). In the Δ*phoPQ* background, the induction of P_*msrA*_::*sfgfp* was highly reduced in intracellular STM (2.61-fold lower x-median RFI).

We conclude that the ability of intracellular STM to manipulate the endosomal system of host cells by action of the SPI2-T3SS effector proteins contributes to reduction of stress exposure, and is especially important in host cells such as phagocytes with a complex set of antimicrobial defense mechanisms.

### *Analyses of the proliferation rate of intracellular* Salmonella *using dual color reporters*

The dual color reporters contain in addition to the stress-inducible sfGFP fusion the constitutively expressed DsRed fusion. The red fluorescence allows detection of STM-infected cells, an estimation of bacterial load per host cell, and the detection of released bacteria in lysates of infected host cells. The constitutive expression of DsRed in combination with the long maturation time of this FP allows the analyses of the intracellular proliferation of STM based on fluorescence dilution by cell divisions (see **Fig. 1**). We used STM WT [P_EM7_::*dsred* P_*msrA*_::*sfgfp*] for population analyses of intracellular STM in RAW264.7 (**Fig. 6**ABCDE) and quantified sfGFP expression as reporter for stress exposure and DsRed fluorescence as indicator for intracellular proliferation. As expected from previous results, intracellular STM showed a strong induction of the P_*msrA*_::*sfgfp* fusion at 8 h p.i. Closer inspection of the DsRed fluorescence intensities indicated heterogenous levels, and plotting of DsRed vs. sfGFP intensities at low cell densities indicated the presence of two subpopulations. We gated subpopulation 1 and subpopulation 2. Quantification of RFI determined 1.61-fold higher sfGFP x-median RFI (**Fig. 6**DE), and 1.59-fold lower DsRed x-median RFI (**Fig. 6**DE) for subpopulation 1 compared to subpopulation 2.

**Fig. 6.**
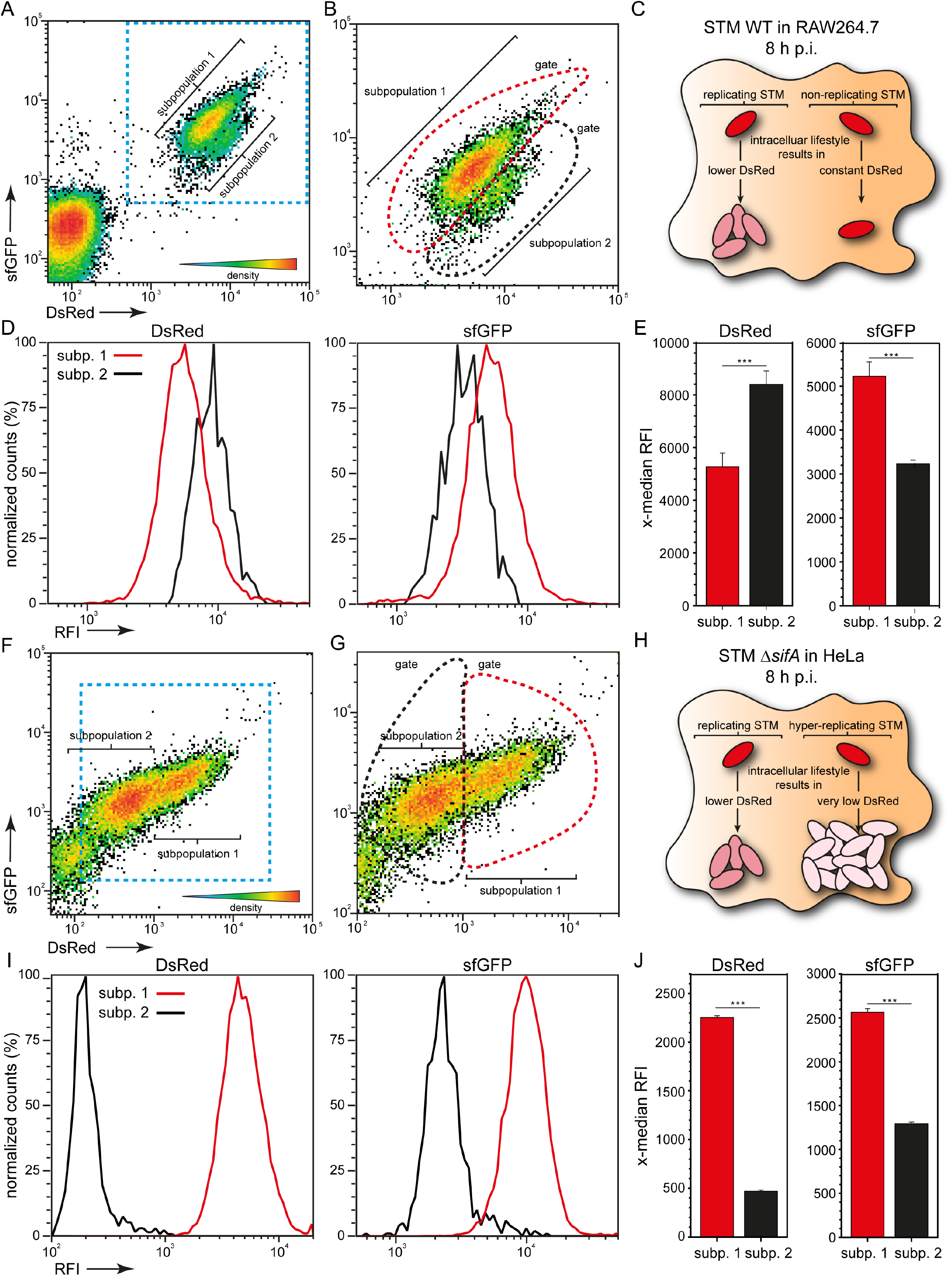
Detection of intracellular subpopulations with distinct stress responses. RAW264.7 macrophages or HeLa cells were infected with STM WT (A-E), or STM Δ*sifA* (F-J), respectively, each harboring dual color fluorescence reporter P_EM7_::*dsred* P_*msrA*_::*sfgfp*. FC analyses of intracellular STM liberated 8 h p.i. from RAW264.7 macrophages (A) or HeLa cells (F). Details are shown in B) and G). By plotting the bacterial population against their constitutive DsRed and P_*msrA*_-induced sfGFP intensity, two subpopulations (subp.) were distinguished. The x-medians represent either constitutive DsRed intensity of intracellular STM, or P_*msrA*_-induced sfGFP signal of DsRed-positive intracellular STM. For WT in RAW264.7 cells the larger subpopulation 1 (red bars) shows lower constitutive DsRed intensity (x-median) compared to the smaller subpopulation 2 (black bars) as indicated in scheme C). Subpopulation 1 shows higher P_*msrA*_-induced sfGFP fluorescence (x-median) compared to subpopulation 2 (D, E). For STM Δ*sifA* in HeLa cells, larger subpopulation 2 mainly consists of cytosolic STM (scheme in panel H) and is characterized by low x-median sfGFP due to lower stress exposure and low x-median DsRed due to rapid growth and DsRed dilution (I, J).

For further test of the reporter system, we investigated the reporter plasmid in STM Δ*sifA* recovered after infection of HeLa cells (**Fig. 6**FGHIJ). This mutant strain shows hyper-replication resulting from fast cytosolic proliferation (see **Fig. 1**H). We were able to distinguish a smaller subpopulation 1 with high DsRed and sfGFP levels, and a larger subpopulation 2 with about 2-fold reduced sfGFP levels and highly reduced x-median RFI for DsRed (**Fig. 6**IJ).

Microscopic analyses confirmed reduced DsRed and sfGFP intensities in STM replicating in the host cell cytosol (data not show). This result indicates that STM in the cytosol of HeLa cells have lower levels of stress exposure and can rapidly replicate.

It should be mentioned that all subpopulations show a high degree of heterogeneity in levels of expression of the stress reporter, and dilution of DsRed. For both FP, about 10-fold differences in intensities can be observed in either subpopulation (**Fig. 6**). We conclude that subpopulation 1 of STM WT in **Fig. 6**AB consists of intracellular proliferating STM with dilution of DsRed. Subpopulation 1 is responding to stress factors in the intracellular environment. In contrast, subpopulation 2 shows reduced intracellular proliferation, resulting in less DsRed dilution and higher levels per cells (**Fig. 6**C). The response to intracellular stress in subpopulation 2 is reduced (**Fig. 6**E).

With the dual color fluorescence reporters, we have at hand a system that allows single cell analyses of response of intracellular STM to environmental stimuli, as well as the interrogation of the proliferation state of the pathogen.

## Discussion

### A versatile system for stress response on single cell level

In this study, we developed, evaluated and applied FP-based reporters for the analyses of the stress response of intracellular STM on a single cell level. We made use of the distinct maturation times of FPs, allowing interrogation of the physiological status of individual bacteria in large and heterogeneous populations. Use of a fast-maturing FP, here sfGFP, allowed sensitive detection of for stress exposure and bacterial response. A second constitutively expressed slow-maturing FP, here DsRed, allowed estimations of the rate of proliferation, since FP dilution by bacterial division occurred more rapidly than synthesis and maturation of new copies of the FP. While we have applied the technique to analyses of STM, the approach will be applicable to other bacteria. This will require selection of a FP with slow maturing kinetics that matches the generation time of the species or strain to be analyzed. Systematic analyses of maturation kinetics, as well as FP half-life and toxicity to the bacterial host will be an important requirement for a rational design of reporters. Such characteristics are partially retrievable from databases such as FPbase (24), but application of FP in a given organism may require direct comparison of FP properties as described here.

While a dual fluorescence approach was used in this study, extension towards a third FP as reporter for additional physiological conditions may be considered. Here, a restriction is the limited separation of FP excitation and emission spectra that may be overcome by improved FC instrumentation and gating approaches. It also has to be considered that FP synthesis levies additional metabolic burden on producing bacteria, and FP can be toxic to the producing cell to various extent. One example is mCherry and its detoxified variant mCherry2 (25). While metabolic burden and cell toxicity are critical parameters in all kinds of infections assays, the importance is even more pronounced in animal models of infection, the generally impose a broader array of stress factors on microbial pathogens resulting in stronger counter-selection.

The criteria for application of FP-based reporter for *S. enterica* in more complex settings such as host tissues have recently been reviewed (26).

Alternatives to FP may be self-labeling enzymes (SLE) that can serve as reporters for promoter activity and, if fused as tags to proteins of interest, as reporters for protein localization and fate (27–29). SLE tags are rapidly maturing reporters and the choice of HaloTag, SNAP-tag and CLIP-tag and a range of fluorochrome-conjugated substrates would allow versatile multiplexing, and combinations with FP. A limitation in use of SLE reporters is the requirement of reaction with cognate substrates and washing to remove unbound substrates.

### Stress response of intracellular STM is dependent on host cell type

We have generated a set of reporters for cytosolic and cytoplasmic SRS in STM. The stress reporters were induced under *in vitro* culture conditions using defined stressors. The reporter response *in vitro* correlated with the dose of the stressor applied, thus providing the option to gauge stressor levels *in vivo*. The long half-life of sfGFP did not allow to analyze potential decrease of stress signals acting on intracellular STM. The addition of tags for degradation, such as the LVA tag (30) is possible and allows generation of dual color reporters with increased dynamic range (J. Röder and M.H., unpublished observations). We observed that induction of all reporters in intracellular STM was higher compared to *in vitro* exposure to defined stressors. This indicates that multiple factors within infected host cells may induce responses of STM in a synergistic manner. The complexity of defense mechanism imposed by host cells thus may restrict the precise quantification of specific levels of single stressors.

As anticipated from the characterization of detoxification or repair systems such as SodAB or DksA, respectively, mutant strains deficient in these systems showed an increased induction of stress reporters. The same effect was observed for mutant strains deficient in the SPI2-T3SS (Δ*ssaV*), or SPI2-T3SS effector proteins required for the efficient remodeling of the host cell endosomal system (Δ*sifA*, Δ*sseF*). Higher stress exposure, especially of SPI2-T3SS-deficient STM was deduced from recent ensemble-based proteomics of intracellular STM (13), and this work confirms the findings with the ultimate resolution of single cell level. The role of SPI2-T3SS for protection of intracellular STM against ROS or RNS has been previous reported (31, 32), suggesting that SPI2-T3SS-mediated redirection of antimicrobial effector mechanisms from the SCV results in protection. More recent work indicated a role of the SPI2-T3SS-mediated formation of an extensive SCV/SIF continuum in nutrient supply to STM within the SCV (12). A potential role of SIF formation in dampening stress by dilution of antimicrobial factors in the lumen of the SCV/SIF continuum has been proposed (11, 33). The single cell analyses shown here support the role of the SPI2-T3SS in protection of intracellular STM against host cell-imposed stressors.

Having established single cell-based stress reporters in STM, it will now be of interest to compare SRS in *S. enterica* serovars to broad host range to serovars that are specialized to infection of human and primates, such as Typhi and Paratyphi A. Such analyses may reveal distinct intracellular adaptation traits as part of the virulence strategy of ‘stealth pathogens’ *S. enterica* serovar Typhi and Paratyphi A (34).

### Proliferating and non-proliferating subpopulations respond to host-imposed stress

The quantification of RFI of two FP in our work allowed to define distinct subpopulations of intracellular STM, showing different levels of intracellular proliferation, and variable degree of stress response reactions. Within the two subpopulations, we defined for STM WT-infected macrophages a smaller subpopulation that showed lower DsRed dilution indicating reduced intracellular proliferation. The induction of the stress response reporter was slightly reduced compared to the more rapidly dividing subpopulation. We conclude that intracellular STM can sense and respond to host cell-imposed stressors, despite residing in a growth-restricted intracellular environment. Thus, mounting efficient stress responses is important for intracellular proliferation of STM, as well as for survival of host defenses in absence of proliferation. This conclusion is also supported by the reduced stress response induction observed for the *sifA* mutant strain. A higher number of bacteria proliferate in the host cell cytosol, and these bacteria encounter a lower level of exposure to stressor.

Our data indicate that stress sensing and stress response is possible in subpopulations with high levels of intracellular proliferation, as well as in cells with reduced rates of proliferation. Next, it will be of interest to analyze the ability in stress response of non-replicating subpopulations. Such subpopulations are also formed by STM in infected macrophages (35), and in infected tissues of host organisms (36). Of particular interest are persister bacteria, since such cells are refractory to most antibiotic treatments, leading the failure of antimicrobial treatment and relapse of infectious diseases (reviewed in 37). The possibility to characterize subpopulations on a single cell level will also enable further studies, such as analyses of stress sensing and response of intracellular persisters. Prior work indicted that STM persisters can actively utilize the SPI2-T3SS to interfere with macrophage polarization by effector protein SteE (38). The expression of genes encoding the SPI2-T3SS is triggered by host stimuli senses by intracellular STM. It will be of interest for further research to identify host stimuli such as stressors that act on the expression of the virulence factor for manipulation of host cells. The dual color reporter approach in combination with FC analyses will be instrumental to detect such events, and should provide access to the very small subpopulation of persister cells. A future perspective is the recovery of distinct subpopulations of intracellular *Salmonella* by fluorescence-assisted cells sorting and global analyses by transcriptomic or proteomic approaches.

## Acknowledgements

This work was supported by the DFG though grants in SFB 944, projects P4 and P15. We kindly acknowledge intramural funding by profile line P2: Integrated Science of the University Osnabrück. We like to thank the Hans-Mühlenhoff-Stiftung for support of KO.

## Materials and Methods

### Generation of reporter plasmids

For the generation of dual color fluorescence reporter plasmids, p4889 (P_EM7_::*dsred* P_*uhpT*_::*sfgfp*) with a constitutive expression of DsRed and GOI-regulated expression of sfGFP was used as described before (Noster *et al.*, 2019). Via Gibson Assembly (GA) of PCR fragments, the *uhpT* promoter was replaced by promoter fragments of *htrA, fkpA, bisC* and *grxA.* Primers are listed in **Table S 1**, and the resulting dual color reporter plasmids are listed in Table 2.

### Generation of complementation plasmid

For the generation of a complementation plasmid, low-copy vector pWSK29 was used as a vector backbone. By GA, the promoter, open reading frame and terminator of the *dksA* gene was inserted as one fragment into pWSK29. Primers are listed in **Table S 1**, and the resulting complementation plasmid is listed in Table 2.

### Generation of deletion mutants

Mutant strains MvP2553 (Δ*dksA*::*aph*), MvP2600 (Δ*dksA*::FRT), and MvP2611 (Δ*phoPQ*::FRT) were generated by λ Red-mediated mutagenesis (39) and mutant allele was transferred into fresh strain background by P22 transduction. Transductants were purified twice on LB agar containing 10 mM EGTA to prevent formation of lysogenic strains. Lysogeny was controlled by streaking onto green plates. In addition, positive clones were confirmed by PCR. The *aph* resistance cassette was removed by FLP-mediated recombination as described before (40). Primers required for generation of deletion, removal of resistance cassette and control of correct insertion are listed in **Table S 1**.

### Bacterial strains and growth conditions

*Salmonella enterica* sv. Typhimurium strain NCTC12023 (STM) was used as wild-type strain and isogenic mutant strains used in this study are listed in Table 1. Bacteria were cultured in lysogeny broth (LB) at 37 °C overnight (o/n) or PCN minimal medium as described in (40), supplemented with 1 mM PO_4_^−^ and 0.4 % glucose or maltose, with a pH of 7.4 at 37 °C using a roller drum at 60 rpm with aeration. For maintenance of plasmids, carbenicillin was added at 50 μg × ml^−1^ as a selection marker.

**Table 1.**
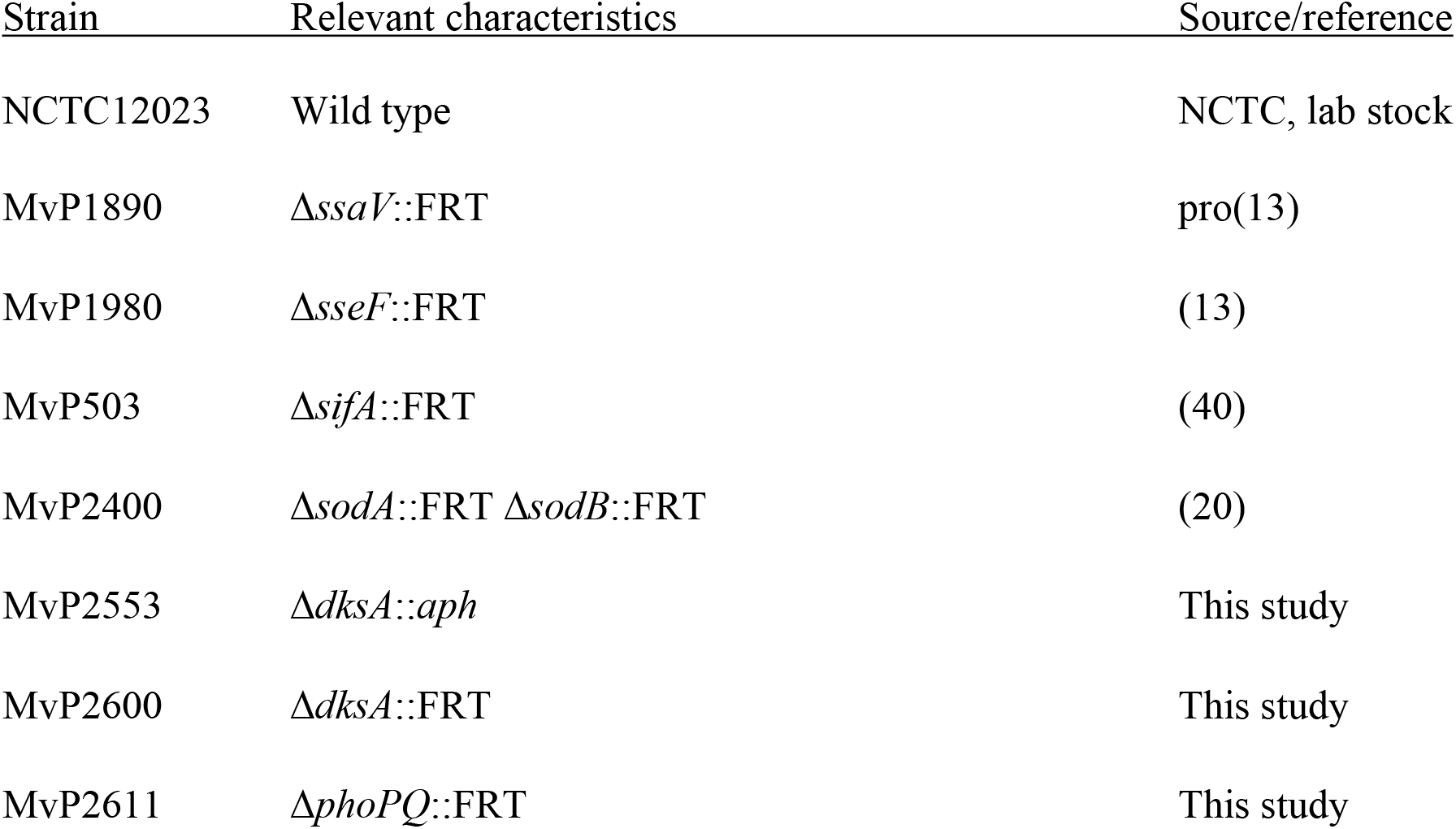
*Salmonella enterica serovar* Typhimurium strains used in this study

**Table 2.** Plasmids used in this study

| 672 | Plasmid | Relevant genotype* | Source |
| --- | --- | --- | --- |
| 673 | p5204 | P<sub>EM7</sub>::dsred P<sub>cypD</sub>::sfgfp (frameshift) | (13) |
| 674 | pFPV-mCherry | P<sub>rpsM</sub>::mCherry | (41) |
| 675 | pWRG167 | P<sub>EM7</sub>::sfgfp | (42) |
| 676 | p5383 | P<sub>EM7</sub>::mCherry2 | This study |
| 677 | p4928 | P<sub>EM7</sub>::tagrfp-T P<sub>tetA</sub>::sfgfp | This study |
| 678 | p5024 | P<sub>dksA</sub>::dksA | This study |
| 679 | p5055 | P<sub>EM7</sub>::dsred P<sub>htrA</sub>::sfgfp | This study |
| 680 | p5084 | P<sub>EM7</sub>::dsred P<sub>msrA</sub>::sfgfp | (13) |
| 681 | p5054 | P<sub>EM7</sub>::dsred P<sub>fkpA</sub>::sfgfp | This study |
| 682 | p5072 | P<sub>EM7</sub>::dsred P<sub>bisC</sub>::sfgfp | This study |
| 683 | p5074 | P<sub>EM7</sub>::dsred P<sub>treA</sub>::sfgfp | (13) |
| 684 | p5085 | $P_{EM7}::dsred\ P_{trxA}::sfGFP$ | (13) |
| 685 | p5086 | $P_{EM7}::dsred\ P_{grxA}::sfGFP$ | This study |
| 686 | * all plasmids confer resistance to carbenicillin |  |  |
| 687 |  |  |  |

### Stress induction by methyl viologen (‘paraquat’), hydrogen peroxide, DTT, polymyxin B, sucrose or heat

For *in vitro* induction of stress, STM strains were cultured overnight in PCN minimal medium (containing 1 mM PO_4_^−^ and 0.4 % glucose), diluted 1:100 in fresh PCN medium and subcultured for further 3 h. Afterwards, bacteria were exposed to methyl viologen (Sigma-Aldrich), hydrogen peroxide (Merck), dithiothreitol, polymyxin B, sucrose or heat to a final concentration as indicated for 3 h at 37 °C with aeration. Subsequently, samples were taken, diluted with PBS and fixed with 3 % *para*-formaldehyde (PFA) in PBS for 15 min at RT. Then, PFA was removed by centrifugation for 5 min at 20,000 *x g* and the pellet was resuspended in 250 μl 100 mM NH_4_Cl in PBS for quenching of residual free aldehydes. After that, samples were directly subjected to FC.

### Cell lines and cell culture

For infection experiments murine RAW264.7 macrophages (American Type Culture Collection, ATCC no. TIB-71) or the non-polarized epithelial cell line HeLa (American Type Culture Collection, ATCC no. CCL-2) was used. RAW264.7 macrophages were cultured in Dulbecco’s modified Eagle’s medium (DMEM) containing 3.7 g × l^−1^ NaHCO_3_, 4.5 g × l^−1^ glucose, 4 mM stable glutamine without sodium pyruvate (Biochrom) and supplemented with 6 %inactivated fetal calf serum (iFCS) (Sigma-Aldrich). HeLa cells were cultured in DMEM containing 4.5 g × l^−1^ glucose, 4 mM stable glutamine and sodium pyruvate and supplemented with 10 % iFCS. All cells were maintained at 37 °C in an atmosphere of 5 % CO_2_ and 90 % humidity.

### Host cell infection for cytometry

Before infection, RAW264.7 macrophages were seeded in surface-treated 6-well plates (TPP) to reach confluency (ca. 2 × 10^6^ cells per well) on the day of infection. If indicated, cells were induced by 2.5 ng × ml^−1^ γ-Interferon at least 24 h before infection. For infection of RAW264.7 macrophages, *Salmonella* strains were grown o/n in LB broth (app. 18 h). Infection was performed with a multiplicity of infection (MOI) of 10. Bacteria were centrifuged onto the cells for 5 min at 500 *x g* and infection proceeded for 25 min at 37 °C in an atmosphere of 5 % CO_2_ and 90 % humidity. Afterwards, infected cells were washed thrice with PBS and incubated for 1 h with cell culture medium containing 100 μg × ml^−1^ gentamicin (Applichem) to kill non-phagocytosed bacteria. Afterwards, the cell culture medium was replaced by medium containing 10 μg × ml^−1^ gentamicin until the end of the experiment. Before infection of Hela cells, they were seeded in surface-treated 6-well plates to reach confluency (~ 1 × 10^6^ cells per well) on the day of infection. For infection of HeLa cells, *Salmonella* strains were grown overnight in LB broth, diluted 1:31 in fresh LB and subcultured for further 3.5 h to induce maximal invasiveness. Infection was performed with a MOI of 5 for 25 min at 37 °C in an atmosphere of 5 % CO_2_ and 90 % humidity. Subsequently, cells were washed thrice with PBS and incubated for 1 h with medium containing 100 μg × ml^−1^ gentamicin (Applichem) to kill non-invaded bacteria. Subsequently, the medium was replaced by medium containing 10 μg × ml^−1^ gentamicin until the end of the experiment.

### Host cell infection for competitive index assays

Before infection, RAW264.7 macrophages were seeded in surface-treated 96-well plates (TPP) to reach confluency (ca. 7 × 10^4^ cells per well) on the day of infection. For infection of RAW264.7 macrophages, STM WT and MvP2553 or STM WT and MvP2600 [p5024] were coinfected with a MOI of 1. Culturing of bacteria and infection conditions were the same as described above. At 2 h post infection (p.i.) and 16 h p.i., host cells were washed twice with PBS and lysed with 0.1 % Triton X-100 in PBS for 10 min at RT with shaking. The lysate was transferred to a test tube, serial dilutions in PBS were prepared and plated in parallel onto LB and LB plates with carbenicillin or kanamycin, respectively. After incubation o/n of plates at 37 °C, the amounts of bacteria per ml after 2 h and 16 h p.i. was determined and the x-fold replication of both strains in the coinfection was calculated. The competitive index or the relative attenuation represents the ratio between both strains in the coinfection.

### Flow cytometry analysis

Flow cytometry (FC) of STM liberated from host cells was performed as described before (Noster et al., 2019). Briefly, PC was performed on an Attune NxT instrument (Thermo Fischer Scientific) at a flow rate of 25 μl × min^−1^. At least 10,000 bacteria were gated by virtue of the constitutive FP fluorescence. Per gated STM cell, the intensity of the sfGFP fluorescence was determined and x-medians for sfGFP intensities were calculated. If required for analyses of STM proliferation, the x-medians of DsRed intensities were determined.

To release the intracellular bacterial population, infected cells were lysed at the indicated time points using 0.5 % Triton X-100 in PBS for 10 min at RT with shaking. The lysate was transferred to a test tube, and after pelleting of host cell debris by centrifugation for 5 min at 500 *x g*, bacteria were recovered from supernatant. Bacteria were further centrifuged for 5 min at 20,000 *x g* and fixed in 3 % PFA in PBS for 15 min at RT. After fixation and a further centrifugation step, fixed bacteria were resuspended in 250 μl 100 mM NH_4_Cl in PBS for quenching of residual free aldehydes. After that, samples were directly subjected to FC. To measure the pre-induction of dual color reporters, bacteria from the inoculum were directly harvested by centrifugation and fixed as described above.

To measure intensity decrease of red fluorescence during growth of a subculture of STM WT within LB or PCN medium supplemented with glucose or maltose or to measure the impact of the maturation time of various red fluorescent proteins on the intensity decrease during growth, STM WT harboring the respective plasmids was grown o/n in the respective growth medium, diluted to an OD_600_ of 0.05 in fresh medium and subcultured over time. Every hour the OD_600_ was determined and samples were taken, diluted in PBS and directly subjected to FC.

## Suppl. Materials

**Table S 1. Oligonucleotides used in this study.**

## Suppl. Figure Legends

**Fig. S 1. Impact of replication status of STM and maturation time of red/green FP on constitutive fluorescence intensity.** A) STM WT harboring plasmids with constitutive expression of either *dsred*, *sfgfp*, tag-*rfpT*, mCherry or mCherry2 was grown o/n in LB medium, diluted to an OD_600_ of 0.05 in fresh LB and incubated at 37°C. Every hour, samples were taken and subjected to flow cytometry (FC) to determine the constitutive fluorescent events per ml (bacteria × ml^−1^) B) STM WT harboring plasmid with constitutive expression of *dsred* was cultured as described above in LB medium. FC measurements were performed as described above to measure the constitutive DsRed intensity. In addition, the OD_600_ was determined. DsRed intensity course shows rapid decrease of DsRed intensity during logarithmic growth phase and recovery of intensity after entering stationery phase. C) STM WT harboring plasmid with constitutive expression of *dsred* was cultured as described above in various growth media. OD_600_ and FC measurements were performed as described above. D) DsRed intensity course shows higher decrease of red fluorescence intensity during logarithmic growth phase when STM shows faster generation time. After entering stationary phase, red fluorescence intensity of STM recovers independently of medium used.

**Fig. S 2. Stress reporter induction using various concentrations of stressors.** A) STM WT harboring dual color fluorescence reporter for *msrA* was grown in PCN minimal medium o/n and diluted in fresh PCN medium for further subculture. After 3 h of subculture, bacteria were stressed for 3 h using various concentrations of methyl viologen or H_2_O_2_ as indicated. Subsequently, samples were taken, fixed and subjected to FC. The x-median represents the P_*msrA*_-induced sfGFP signal of the entire DsRed-positive bacterial population (left panel), and histograms show population analyses of response to H_2_O_2_ (middle panel), or methyl viologen (right panel). B) STM WT harboring dual color fluorescence reporter for P_*htrA*_ was grown in PCN minimal medium o/n and diluted in fresh PCN medium for further subculture. After 3 h of subculture, bacteria were stressed for 3 h using various concentrations of DTT or polymyxin B as indicated. Analyses was performed as for A), showing x-median of P_*htrA*_-induced sfGFP signal of the entire DsRed-positive bacterial population (left panel), and histograms show population analyses of response to DTT (middle panel), or polymyxin B (right panel). C) STM WT harboring dual color fluorescence reporter for *treA*, *htrA*, or *fkpA* representing the periplasmatic stress response (PSR), and *msrA*, *trxA*, *grxA*, or *bisC* representing the cytoplasmic stress response (CSR) were grown o/n in PCN minimal medium, and diluted in fresh PCN medium for further subculture. At 6 h or 24 h of subculture, samples were taken, fixed and subjected to FC. As a non-induced negative control, the same dual color reporter backbone was used with a frame shift mutation in *sfgfp* (described before in (13). The x-median represents the pre-induced sfGFP signal without stressor addition of the entire DsRed-positive bacterial population (left), and histograms show population analyses of 6 h subcultures (middle panel), or 24 h subcultures (right panel). D) Pre-induction of stress reporter in PCN minimal medium without stressor addition in stationary phase (o/n culture) is shown. Samples were normalized to pre-induction after 6 h subculture.

**Fig. S 3. Time-resolved stress response of dual color fluorescence reporters in STM WT and SPI2-T3SS mutant strains.** STM WT, Δ*sseF*, Δ*sifA* and Δ*ssaV* harboring dual color fluorescence reporter for *msrA* were grown o/n in LB medium. RAW264.7 macrophages were infected using reporter strains and lysed 8 h, 16 h, 24 h and 48 h p.i. Liberated STM were recovered, fixed and subjected to FC. As an uninduced negative control, the same dual color reporter backbone was used with a frame shift mutation in the gene of sfGFP (described before in Noster et al., 2019). The x-median represents the induced sfGFP signal of the entire DsRed-positive intracellular bacterial population (B) and the respective histograms are shown in A). C) The depicted slopes represent the sfGFP intensity increase over the time of infection (calculated of B). Higher stress induction over a constant period of time results in a higher slope as shown for example for Δ*ssaV*. The same was measured for intracellular STM harboring dual color fluorescence reporter for *trxA* and *htrA*. Means and standard deviations of one representative experiment are shown. Statistical analysis was performed as for Fig. 3.

**Fig. S 4. Induction of P_*msrA*_ in deregulated *phoPQ* null mutant 8 h p.i. in RAW264.7 macrophages.** STM WT and Δ*phoPQ* strains harboring dual color fluorescence reporter for *msrA* were grown o/n in LB medium. RAW264.7 macrophages were infected and lysed 8 h p.i. Liberated STM were recovered, fixed and subjected to FC. As an uninduced negative control, the same dual color reporter backbone was used with a frame shift mutation in *sfgfp*. The x-median represents the P_*msrA*_-induced sfGFP signal of the entire DsRed-positive intracellular bacterial population in A), and a representative histogram is shown in B). Means and standard deviations of one representative experiment are shown.

